# CaNetiCs - An Open-Source Toolbox for Standardized Dimensionality Reduction of Neuronal Calcium Activity

**DOI:** 10.1101/2025.08.01.668030

**Authors:** Daniel Carbonero, Jad Noueihed, Christopher V. Gabel, Mark A. Kramer, John A. White

## Abstract

The widespread use of calcium imaging has produced large-scale datasets capturing neuronal population activity across diverse experimental contexts, posing challenges for analyzing complex, high-dimensional data. Dimensionality reduction (DR) methods have been pivotal in addressing these challenges by simplifying data into interpretable, low-dimensional structures, while capturing essential network dynamics. Among DR methods, Nonnegative Matrix Factorization (NMF) can produce biologically meaningful representations through its nonnegativity constraint and parts-based decomposition, making it especially suited for analyzing neuronal calcium signals. To enhance accessibility and standardization in the analysis of state-dependent neuronal dynamics, we introduce **Ca**lcium **Net**work dynami**Cs** (CaNetiCs), an open-source toolbox centered on NMF, integrating standardized DR methods (PCA, ICA, UMAP), geometric low-dimensional component space analyses, and neuronal network simulation modules.

We validate our toolbox by applying it to two diverse experimental datasets that describe responses to graded anesthesia: whole-ganglion cellular calcium imaging of C. elegans and two-photon imaging of murine somatosensory cortex. Our analyses recapitulate previously observed trends, such as network suppression and decorrelation with anesthesia, while uncovering novel insights into neuronal activity under differing contexts. CaNetiCs provides an accessible, modular, and interpretable framework, facilitating broader adoption of standardized dimensionality reduction methodologies for deeper exploration of neuronal network dynamics across experimental paradigms. The open-source code, along with documentation, is available at https://github.com/dannycarbonero/CaNetiCs.

## Introduction

The ability to record activity from large populations of neurons with single-cell resolution has revolutionized neuroscience, enabling new insights into how distributed networks dynamically encode behavioral and physiological states^1^. Calcium imaging, in particular, allows monitoring of hundreds to thousands of neurons simultaneously across different brain regions and experimental conditions^2–6^. However, the complexity and dimensionality of these recordings pose significant challenges for analysis. Standard approaches, such as computing descriptive statistics or averaging across cells and time points, often obscure the rich temporal and population-level structures inherent in the data^7,8^.

Dimensionality reduction (DR) techniques provide a powerful solution by projecting high-dimensional neural recordings onto lower-dimensional representations, retaining major sources of variance while facilitating interpretation of collective dynamics^9^. Linear DR methods, such as Principal Component Analysis (PCA)^10–16^ and Independent Component Analysis (ICA)^17,18^, have been widely used to identify patterns of coordinated neuronal activity. More recently, nonlinear methods such as Uniform Manifold Approximation and Projection (UMAP) have been employed to capture complex structures in neural population data^19^. Yet, each approach imposes distinct mathematical assumptions, such as orthogonality in PCA, independence in ICA, or manifold continuity in UMAP^20–22^, that can influence the nature and interpretability of the extracted components.

Nonnegative Matrix Factorization (NMF) represents a particularly promising alternative for calcium imaging analysis^15,18^. By constraining both the data and its decomposition to be nonnegative^23^, NMF preserves the additive nature of fluorescence signals and produces parts-based representations, wherein each component corresponds to an interpretable sub-network of activity^15,18^. Unlike PCA or ICA, which impose strict mathematical constraints to construct a less interpretable low-dimensional representation, NMF decomposes neuronal activity into coherent modules that can be directly associated with underlying biological processes^18^. Prior work has demonstrated that NMF robustly captures dominant organizational features of neuronal activity in simulated networks, and outperforms alternative DR methods in reconstructing known ground-truth structures^18^.

Although dimensionality reduction methods offer powerful means for interpreting neuronal population dynamics, their application often requires specialized technical knowledge and complex, customized computational workflows. Sophisticated analyses, including variance optimization, information-theoretic model selection, and geometric modeling of state spaces, can be difficult to implement without significant computational expertise. Recent work has led to an increase of open-source software toolboxes emerging to lower the barrier to entry for complex data analyses across fields of neuroscience^15,24,25^. However, available toolboxes remain limited with respect to providing standardized, tunable, and feature-rich user-friendly frameworks specifically tailored for analyzing state-dependent neuronal dynamics from calcium imaging recordings. To address this gap, we developed **Ca**lcium **Net**work dynami**Cs** (CaNetiCs), an integrated toolbox that consolidates validated analytical methods into a flexible and accessible codebase.

Building on our prior work^18^, our software offers a standardized implementation of NMF-based dimensionality reduction optimized for calcium recordings, alongside complementary DR approaches (PCA, ICA, UMAP), geometric low-dimensional component space analysis, simulation tools for method benchmarking, and basic neuronal network simulation analysis, providing modularity for hypothesis-driven and exploratory analyses alike. A key advantage of our NMF decomposition implemented here is the creation of easily interpretable structures: lower-rank components that correspond to distinct patterns of activity, or sub-networks of neuronal activity, and the associated neuronal weights that reflect the extent to which each neuron contributes to each sub-network^15,18,26^. Each neuron’s activity is thus approximated as a positive, additive combination of a small number of dominant modes. This parts-based representation^27^ provides an intuitive map of network organization, identifies leading drivers of population dynamics, and facilitates quantitative comparisons of neuronal sub-network structures across experimental conditions. Moreover, the nonnegativity constraint ensures that all extracted features remain biologically plausible, avoiding artifacts introduced by cancellation effects inherent to other DR methods^18,20^.

We demonstrate the utility of CaNetiCs by building a standard analytical pipeline that applies our DR analysis to two diverse experimental datasets: light-sheet calcium imaging of nearly the entire head ganglia of Caenorhabditis elegans under graded isoflurane anesthesia^16^, and two-photon imaging of murine somatosensory cortex (S1 L2/3) across awake, anesthetized, and recovery states^28^. Across both datasets, we find that CaNetiCs not only recapitulates previously reported macroscopic trends in network suppression and decorrelation with anesthesia but also uncovers novel geometric and dynamic features of state transitions at the population level. By providing an accessible, flexible, and interpretable framework for dimensionality reduction analysis, we seek to enable a broader adoption of systematic and sophisticated network-level approaches for a diversity of calcium recordings under various contexts, and to facilitate finding deeper insights into how large-scale neuronal dynamics are shaped by experimental perturbations, biological state, and species-specific differences.

## Results

We first apply our developed pipeline to analyze calcium recordings under increasing concentrations of isoflurane anesthesia in *C. elegans*^16^ and in mice^28^. We fit models to the optimal dimensionality informed by an information criterion^29–31^ and to three dimensions, consistent with previous analyses^10,13,16,32–35^, to analyze the neuronal network dynamics under differing concentrations of anesthesia. We show that our analyses naturally capture the different dynamics between the species, support the conclusions of previous descriptive analyses, and provide further novel insights to differential dynamics as a function of anesthetic concentration.

### Neuronal network dynamics of C. elegans under increasing concentrations of isoflurane

We analyze the neuronal network dynamics of anesthesia recorded in *C. elegans* under no (0% Isoflurane), intermediate (4% Isoflurane), and deep (8% Isoflurane) anesthesia. We re-use these data previously reported in Awal et al^16^. For each animal (n = 10) we apply the CaNetiCs pipeline to the calcium traces (examples in Fig.1a) as follows.

We first iteratively fit models for 25 components, for each animal, for each condition. Then averaging across animals, we investigate variance explained and information criterion across conditions. We begin by analyzing the variance explained across conditions. We find that variance is explained most quickly in the awake state, followed by the deeply anesthetized state, and then the intermediately anesthetized state (Fig. 1c). Consistent with previously conducted analyses of the same data set^16^, most of the variance explained is captured within three components. Previous analysis of individual trace^16^ activity determined that neurons are significantly more correlated during the awake state, followed by intermediate anesthesia, and deep anesthesia, with the only statistically significant difference being between the awake and anesthetized states. Because AIC follows from variance explained, optimizing between model complexity and variance captured, AIC follows the same trend as variance explained, but inverted. AIC is lowest during the awake state, followed by the deeply anesthetized state, and finally the intermediary anesthetized state (Fig. 1d). We then fit models to the optimum number of components determined by the Akaike Information Criterion (AIC), for each animal, for each state, where the optimal dimensionality is determined as the minimum of the Akaike Information Criterion (AIC) ^29–31^ (Fig. 1d). We find that the inferred number of optimal components (Fig. 1b) across states follows the same correlation structure found when analyzing singular neuronal correlations in [16], with the awake state requiring the lowest dimensionality, followed by the deep and intermediate anesthetized states, respectively. We find the difference between the optimal components to be statistically significant (Wilcoxon signed-rank test, corrected for multiple comparisons, Table 1) between the awake state and the intermediate state, with the difference trending towards significance between 4% and 8% anesthesia. This suggests a dimensionality difference between 4% Isoflurane and the other states, while the awake state and deeply anesthetized state can be explained by similar dimensionality, but with different activity profiles. While the intermediate state of anesthesia displayed an intermediate activity profile, it showed the least correlated dynamics demonstrating that, while activity consistently dropped as a function of deeper anesthesia, correlations dropped but then once-again increased under deeper anesthetic^16^.

**Fig. 1.**
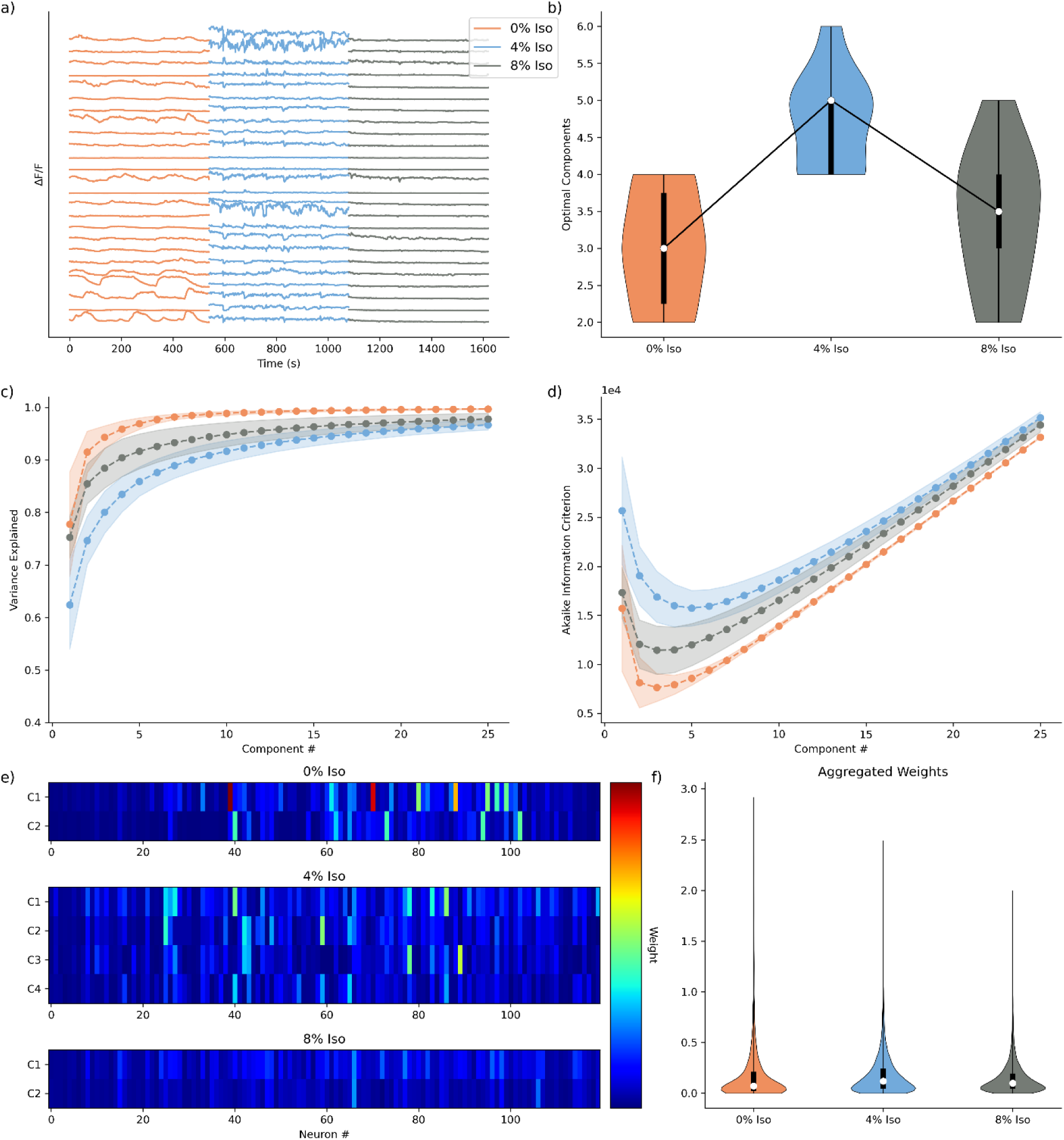
Optimal Dimensionality Models for C. elegans Isoflurane Data Set. **a)** Activity traces for a subset (n=25) of neurons, for a single animal, across conditions. **b)** Distribution of optimal components for all animals, across states (n = 10). **c)** Mean variance explained curve with standard deviation intervals (n=10). **d)** Mean Akaike Information Criterion (AIC) curve with standard deviation intervals (n=10). **e)** Visualization of neuronal weights at optimal dimensionality for animal shown in a), standardized across conditions for comparison. **f)** Aggregated neuronal weights for each animal at optimal dimensionality (n = [3600, 5640, 4080], respectively).

**Table 1.** Significance values for statistical comparison of optimal number of components found in Fig. 1b). Wilcoxon Signed-Rank Test, corrected for multiple comparisons, n = 10.

| Comparison | Components p-value |
| --- | --- |
| 0% Iso - 4% Iso | 0.023 |
| 0% Iso - 8% Iso | 1.000 |
| 4% Iso - 8% Iso | 0.094 |

**Table 2.** Significance values for statistical comparison of volume expanse in space (Fig. 2f). Wilcoxon Rank-Sum Test, corrected for multiple comparisons, n = 10.

| Comparison | Volumes p-value |
| --- | --- |
| 0% Iso - 4% Iso | 0.289 |
| 0% Iso - 8% Iso | 0.004 |
| 4% Iso - 8% Iso | 0.020 |

We next visualize the neuronal weights decomposed under each condition qualitatively for a single animal’s experimental recording (Fig. 1e) and aggregated across animals for all components (Fig. 1f). In addition to requiring fewer dimensions to optimally explain the awake and deeply anesthetized states, compared to the intermediate state, a few neurons are found to drive most of the activity (Fig 1e). Further, normalizing the scale across the decompositions for the different states, we qualitatively find the highest weights in the model for the awake state, especially for the first component (Fig. 1e). Examining the distributions of aggregated weights for all components across states (Fig. 1f), we find the weights distribute similarly to the total macroscopic trends of activity in the data^16^, with the deep anesthetization weights being the most suppressed, the intermediate anesthetization having the widest distribution, and the awake state having the longest tail. These results demonstrate that our analytical pipeline is capturing both the dynamics and the dimensionality of the data in an unsupervised dimensionality reduction model.

Having chosen the number of dimensions, we proceed by fitting models to three dimensions for all experimental states to build a low-dimensional component space model (Fig. 2), building upon previous analytical methods, and furthering previous analysis^10,13,16,32–35^.

**Fig 2.**
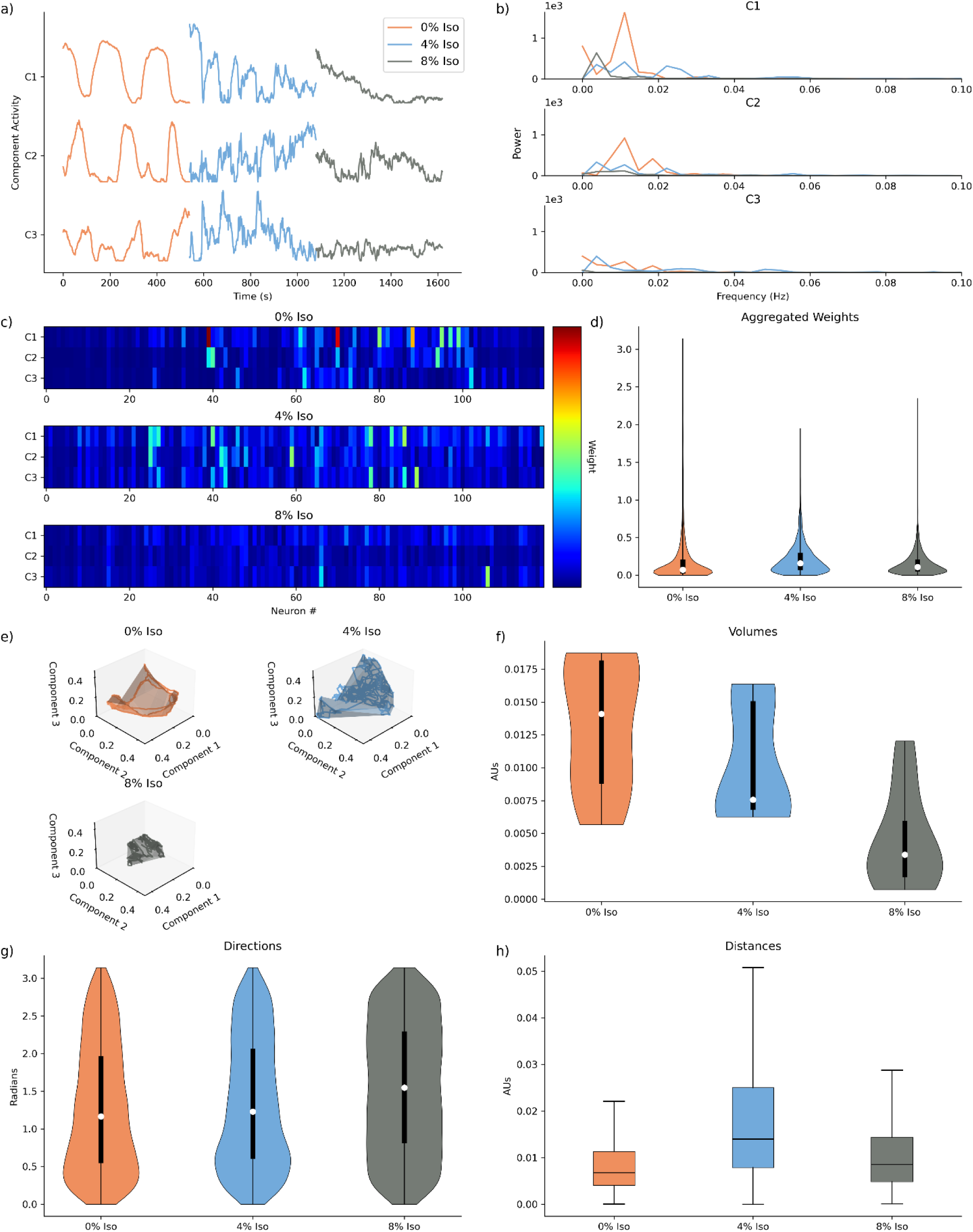
Three-Dimensional Component Space Models for C. elegans Isoflurane Data Set. **a)** Component activity traces for first three components for the same animal shown in Fig. 1a). **b)** Decomposed frequency content for components shown in Fig 2a). **c)** Visualization of neuronal weights for single three-dimensional models for the same animal in Figs. 2a,b). **d)** Aggregated neuronal weights for all three-dimensional models, across animals and states (n = 3600). **e)** Plotted low-dimensional component space of components against each other, with encapsulated volume, for animal shown Fig. 2a,b,c). **f)** Distribution of volumes, across animals and states (n=10). **g)** Distribution of directions, across animals and states (n=5380). **h)** Distributions of distances across animals and states (n = 5380)

We first qualitatively examine neuronal dynamics for a representative animal (the same animal utilized in Fig. 1). By visualizing the activity profiles of the first three components across different anesthetic conditions (0% Isoflurane, 4% Isoflurane, and 8% Isoflurane), we observe clear distinctions that align closely with previous analyses of this dataset^16^. Specifically, the awake state (0% Isoflurane) exhibits a highly structured activity pattern with a distinctly ordered, nearly oscillatory behavior across all three components. In contrast, the intermediate anesthetic state (4% Isoflurane) displays a disrupted and scrambled activity pattern, reflecting a transitional and dynamically unstable neuronal state. The deeply anesthetized state (8% Isoflurane) reveals markedly suppressed neuronal activity, with greatly diminished amplitude across components (Fig. 2a). These qualitative observations corroborate prior findings, offering visual validation of differential neuronal network dynamics across anesthetic states.

To characterize further these oscillatory characteristics, we perform spectral analyses to assess the frequency content within each component across anesthetic conditions. Consistent with our qualitative observations, spectral analysis reveals prominent low-frequency peaks specifically within the awake state, consistent with oscillatory dynamics. These peaks are notably attenuated or absent in both intermediate and deeply anesthetized states, underscoring the disruption and suppression of structured neuronal oscillations as anesthetic depth increases (Fig. 2b).

We next visualize the neuronal weights derived from a three-dimensional decomposition, again for the same representative animal. Qualitatively, we observe a trend consistent with our earlier findings from optimal dimensionality analysis (Fig. 1); neuronal activity in the awake state is driven by larger neuronal weights compared to anesthetized conditions (Fig. 2c). Quantitative analysis of neuronal weights across the full dataset further supports these observations. We detect a similar overall relationship to that observed with optimal component decompositions, though with a notable difference: the intermediate anesthetic state’s distribution tail does not exceed that of the deeply anesthetized state. This discrepancy arises from truncating the weight distribution due to the fixed three-dimensional decomposition rather than utilizing state-specific optimal dimensionalities (Fig. 2d).

To further illustrate the dynamics, we plot the dynamics projected to these first three components in three-dimensional space. We find a distinctly ordered movement about this space in the awake state, reflective of the pronounced oscillatory structure. In contrast, the intermediate state displays less well-ordered dynamics, and the deeply anesthetized state shows marked suppression and minimal spatial exploration (Fig. 2e). To characterize the extent of these different dynamics, we compute the volume fit to the curve across all animals and observe the largest volumes in the awake state, followed by intermediate, and the smallest in the deep anesthetic state; the deep state significantly differs from both awake and intermediate states (Wilcoxon Rank sum test, n = 10, corrected for multiple comparisons). These results align with the observed magnitude of neuronal activity differences across conditions (Fig. 2f).

Analyzing the directional trajectories of the movement about space, we find highly consistent, smooth directional changes in the awake state, a marginal decrease in smoothness in the intermediate state, and slightly more irregular directional changes in the deep anesthetic state (Fig. 2g). Finally, examining the distances traveled through the low dimensional space, we find that the awake state traverses the shortest distances per unit time, followed by the deep anesthetic state, and then the intermediate anesthetic state (Fig. 2h). These patterns of a smoother, slower, movement in state space are consistent with the previously identified higher neuronal correlation profiles reported in Awal et al^16^.

Overall, we conclude the awake state’s movement is more expansive (i.e., covers more of the lower dimensional space) and more directed and smooth compared to the anesthetized conditions. Here, by leveraging both qualitative and quantitative analyses across geometric and temporal domains to create a DR framework for calcium recordings, our approach corroborates earlier findings^16^ and introduces a unified framework for interpreting complex state-dependent network dynamics while adding novel insight.

### Neuronal network dynamics of murine S1 L2/3 under increasing concentrations of isoflurane

As a second illustration of the CaNetiCs pipeline, we analyze calcium recordings from layer 2/3 of murine S1 (n = 36 rodents). The experimental context begins with imaging of an awake or baseline state, increases to an intermediate anesthetic concentration (0.7% Isoflurane), proceeds to a deeper anesthetic concentration (1.4% Isoflurane), and ends with imaging of a recovery state with no anesthetic (examples in Fig. 3a). In Noueihed et al ^28^, the authors used these data to show that single-cell neuronal activity becomes more sparse and more uniform, as a function of entering deeper levels of anesthetic sedation. Here, we re-use these data to illustrate application of the CaNetiCs pipeline.

**Fig. 3.**
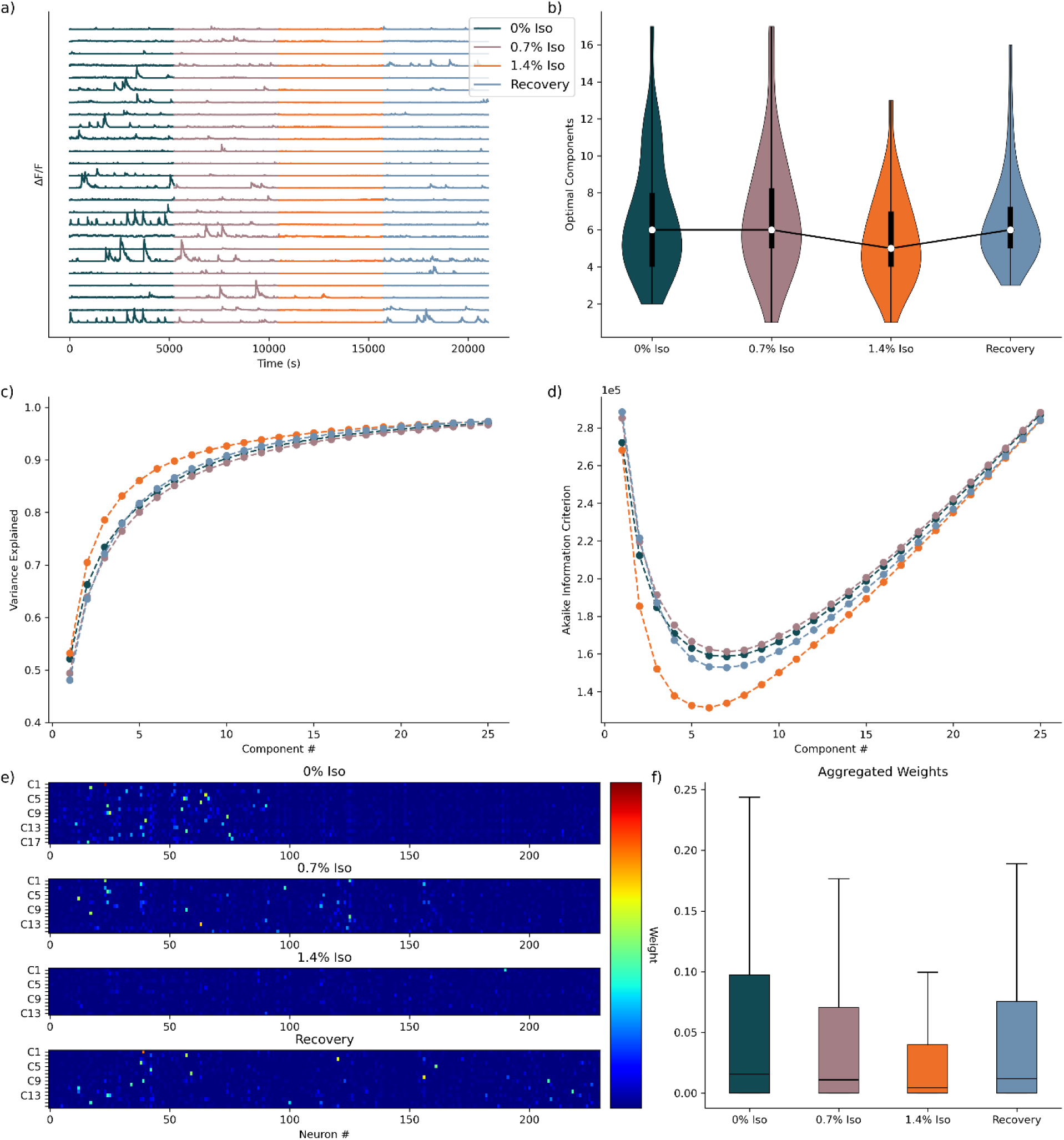
Optimal Dimensionality Models for Murine Isoflurane Data Set. **a)** Activity traces for a subset (n=25) of neurons, for a single animal, across conditions. **b)** Distribution of optimal components for all animals, across states (n = 36). **c)** Mean variance explained curve with standard deviation intervals (n=36). **d)** Mean Akaike Information Criterion (AIC) curve with standard deviation intervals (n=36). **e)** Visualization of neuronal weights at optimal dimensionality for animal shown in a), standardized across conditions for comparison. **f)** Aggregated neuronal weights for each animal at optimal dimensionality (n = [27711, 27589, 22354, 26388], respectively).

We begin by analyzing the variance explained versus the number of components across the three anesthetic conditions. We find that variance was most rapidly captured by the deeply anesthetized state (1.4% Isoflurane), with a distinct gap between this condition and the others (Fig. 3c). This finding aligns closely with prior analyses of single neuronal activities, which demonstrated a marked decrease and increased uniformity or correlation of neuronal activity under deep anesthesia^28^. The variance explained analysis corroborates these findings by rapidly capturing the simplified, more correlated neuronal activity patterns characteristic of deep anesthesia. Computing the Akaike Information Criterion (AIC), we observe a similar trend (Fig. 3d). The deep anesthetic state consistently showed the lowest AIC values, indicating lower model complexity across components due to the relatively simpler activity profile. Evaluating the optimal number of components inferred from the minimum of the AIC curves, we find that the deeply anesthetized state (1.4% Isoflurane) requires significantly fewer components to represent neuronal activity compared to all other anesthetic conditions (Table 3, Wilcoxon Signed-Rank test, corrected for multiple comparisons, n=36). No significant differences emerged among the other anesthetic conditions regarding optimal dimensionality (Fig. 3b).

**Table 3.**
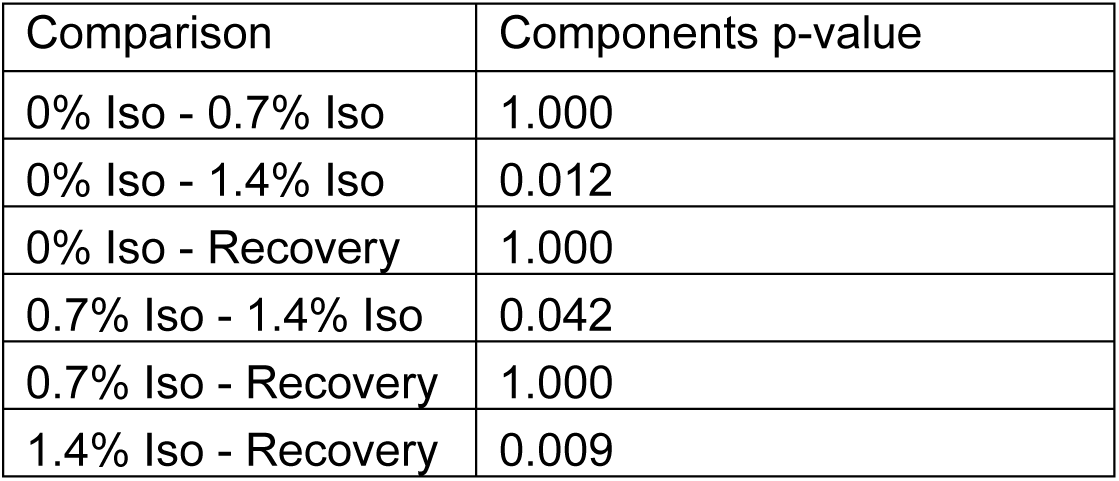
Significance values for statistical comparison of optimal number of components found in Fig. 3b). Wilcoxon Signed-Rank Test, corrected for multiple comparisons, n = 36.

Examining the neuronal weights for a model representing a single animal recording, we find results consistent with previous analyses. Specifically, visual inspection (Fig. 3e) suggests neuronal weights are consistently highest during the awake state, indicating active neuronal contributions. Intermediate and recovery states exhibited more moderate neuronal contributing to the model, but with increased sparsity, with fewer neurons reaching this moderate level of contribution. Finally, the deeply anesthetized state demonstrated comparatively minimal neuronal weights. To characterize these observations, we aggregate neuronal weights across all animals for each experimental condition and find neuronal weights are highest in the awake condition, followed sequentially by the recovery and intermediate states, with the deeply anesthetized state consistently demonstrating the lowest neuronal weights (Fig. 3f).

Taken together, these results highlight the capacity of our DR approach to robustly capture the underlying structure of neuronal activity in reduced-dimensional representations, adjusting dimensionality as the anesthetic conditions informed the information criterion. Importantly, our analyses consistently suggest the presence of an intermediate state, distinct from both the deeply anesthetized and awake baseline states. This intermediate state is characterized by moderate neuronal activity complexity, driven by lower component weights and reduced interactions between neuronal networks, along with decreased overall neuronal activity. However, it maintains a dimensionality similar to that observed in the awake state. This intermediate state, partially uncovered here and suggested by previous studies^28^ emphasizes the nuanced neuronal dynamics underlying anesthetic-induced changes in cortical activity. Finally, we fit three-dimensional models to each animal across all experimental states to further analyze the neuronal dynamics. Visual inspection reveals activity patterns consistent with previous analyses of the neuronal activity^28^ The awake state displays the most extensive activity, followed by intermediate activity levels during intermediate anesthesia (0.7% Isoflurane) and the recovery state, with significantly reduced activity evident in the deeply anesthetized state (1.4% Isoflurane) (Fig. 4a). Spectral analyses revealed the awake state potentially trending towards higher frequency activity than any other state, but the relationship is not robustly apparent (Fig. 4b).

**Fig 4.**
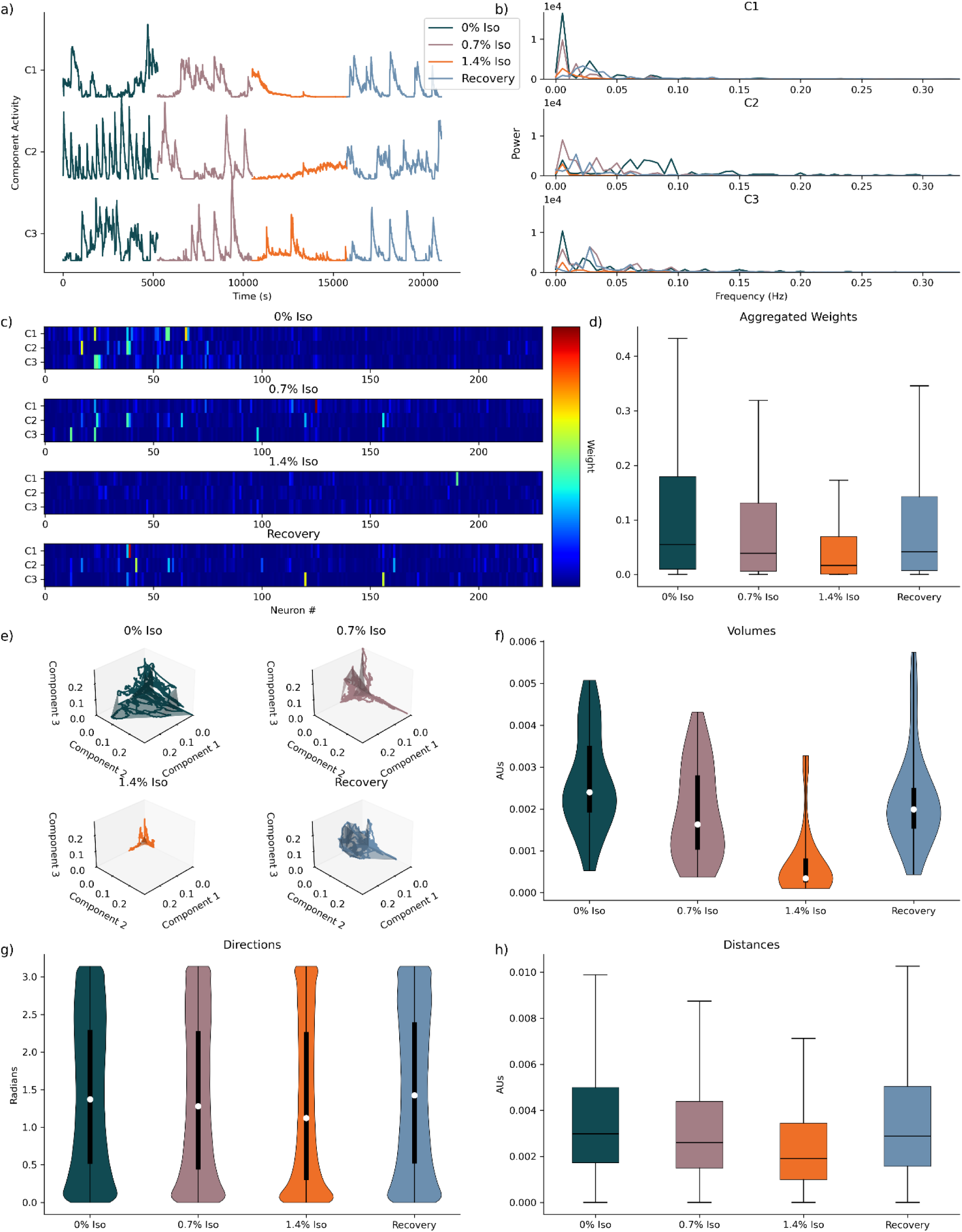
Three-Dimensional Component Space Models for Murine Isoflurane Data Set. **a)** Component activity traces for first three components for the same animal shown in Fig. 1a). **b)** Decomposed frequency content for components shown in Fig 2a). **c)** Visualization of neuronal weights for single three-dimensional models for the same animal in Figs. 2a,b). **d)** Aggregated neuronal weights for all three-dimensional models, across animals and states (n = 10743). **e)** Plotted low-dimensional component space of components against each other, with encapsulated volume, for animal shown Fig. 2a,b,c). **f)** Distribution of volumes, across animals and states (n=36). **g)** Distribution of directions, across animals and states (n=188928). **h)** Distributions of distances across animals and states (n = 188928)

Qualitative visualizations, of neuronal weights decomposed into three components for a single mouse recording, indicate trends consistent with the more comprehensive optimal models, exhibiting increased sparsity, with fewer neurons making meaningful contributions. This sparsity likely reflects the inherent limitation of capturing the full complexity of neuronal dynamics within just three dimensions, underscoring the need for higher-dimensional representations to fully encapsulate the data (Fig. 4c). Expanding this analysis quantitatively by aggregating neuronal weights across all animals confirmed the observed qualitative trends. The awake state again showed the highest neuronal weights, with intermediate and recovery states presenting moderate weights, and the deeply anesthetized state consistently exhibiting minimal contributions. This suggests that although reduced dimensionality captures broad dynamics, it misses details present in optimal higher-dimensional models (Fig. 4d).

Further, plotting neuronal activity in three-dimensional state space revealed that the awake state demonstrated the most expansive and dynamic exploration of this space, consistent with its greater and more diverse neuronal activity (Fig. 4e). Quantitative assessments of volumes encapsulating these trajectories showed statistically significant smaller volumes in the deeply anesthetized state compared to all other states (Table 4, Wilcoxon Rank-Sum test, corrected for multiple comparisons, n=36). Moreover, an emerging, though not statistically significant, trend indicated lower spatial volumes for the 0.7% Isoflurane and recovery states versus the 0% Isoflurane state, highlighting potential intermediary characteristics between awake and moderately anesthetized conditions (Fig. 4f).

**Table 4.** Significance values for statistical comparison of volume expanse in space, found in Fig. 4f). Wilcoxon Rank-Sum Test, corrected for multiple comparisons, n = 36.

| Comparison | Components p-value |
| --- | --- |
| 0% Iso - 0.7% Iso | 0.052 |
| 0% Iso - 1.4% Iso | 4.05e-09 |
| 0% Iso - Recovery | 0.360 |
| 0.7% Iso - 1.4% Iso | 1.41e-06 |
| 0.7% Iso - Recovery | 1.000 |
| 1.4% Iso - Recovery | 4.86e-08 |

Analyzing directional movement in the three-dimensional space revealed a shift from predominantly stochastic trajectories with a preference for small directional changes in the awake state, towards more distinct bimodal directionality under deep sedation, further positioning intermediate anesthesia and recovery states as transitional (Fig. 4g). Finally, evaluating distances traveled through the three-dimensional space revealed relatively similar trajectories across all states, excluding the deeply anesthetized state, which consistently showed reduced movement.

Collectively, these analyses reinforce the utility of using a three-dimensional component space framework for capturing key aspects of neuronal dynamics and revealing nuanced insights, such as a potential intermediate state as a function of entering and exiting anesthesia, that might not immediately be evident from single-neuron analyses alone. The relationships observed among states, dimensionalities, activities, and transitional states underscore complex neuronal dynamics modulated by anesthetic depth and recovery.

Overall, our analytical pipeline extracts low-dimensional representations of neuronal calcium recordings that not only recapitulate findings from previous analyses but also enable novel insights into neuronal network dynamics across experimental conditions. The combination of information-theoretic optimization with dimensionality reduction yields interpretable structures that align with known biological phenomena, such as anesthetic-induced suppression and decorrelation of neural activity. Moreover, we demonstrate that our unsupervised approach can robustly uncover markedly different underlying structures across species, highlighting the method’s generalizability and broad applicability. By providing both quantitative and qualitative validation of network dynamics, we demonstrate that our toolbox offers a unified, flexible framework for analyzing diverse neuronal datasets. These results illustrate that dimensionality-reductive modeling, when standardized and optimized as in our toolbox, not only confirms previous findings but also opens new avenues for hypothesis generation and deeper biological interpretation.

## Discussion

Here, we present an integrated analytical pipeline centered around NMF for the dimensionality reductive analysis of neuronal calcium recordings and wrap this framework into an open-source toolbox. Our goal was to provide a standardized, tunable, and interpretable approach for reducing complex neuronal population activity into tractable low-dimensional representation with easily implementable software.

We validated our approach by applying it to two distinct datasets, calcium recordings from C. elegans and L2/3 murine S1 cortex under varying depths of isoflurane anesthesia. Across both datasets, our method robustly captured biologically meaningful differences in neuronal dynamics between experimental states^16,28^. Notably, the low-dimensional structures extracted by NMF closely concurred with previous analyses of these datasets, while providing new avenues for quantitative and geometric analysis, such as characterizing the expanse of network activity in state space and the smoothness of neuronal trajectories moving in a dimensionally reduced space. Importantly, we observed that the extracted low-dimensional representations differed substantially between species, following true underlying biological distinctions, rather than analytical artifacts. This ability to adaptively capture species-specific network structures in an unsupervised manner underscores the generalizability and broad utility of the approach. Further, when comparing three-dimensional decompositions to the optimal dimensionality fits, we observed an important difference between species. In C. elegans, where the underlying neuronal dynamics are comparatively simpler, fitting to three components recapitulated much of the structure captured at the optimally inferred dimensionality. This suggests that the dominant modes of neuronal activity in C. elegans are highly compressible and well-approximated by this lower-dimensionality state space. In contrast, the murine S1 cortex recordings, characterized by more complex and heterogeneous dynamics, revealed that the three-dimensional models began to capture broad trends across anesthetic states but failed to fully encapsulate the richness of the underlying activity. Here, the reduced dimensionality truncated finer structure, particularly in intermediate and recovery states. This highlights a critical consideration. While low-dimensional representations are highly informative, the appropriate dimensionality must be chosen with regard to the biological complexity of the system under study.

Given the hierarchical structure of our data, we focused our statistical analyses on measures aggregated at the level of the animal, such as optimal dimensionality and volumes spanned in state space, rather than on per-point measures like individual neuron weights, distances, or directions, which exist at vastly higher scales (10³–10⁵ points per condition, detailed in the Fig. captions for each distribution). While the modest number of animals naturally limits the statistical resolution attainable for detecting subtle effects, our aim was not to derive absolute metrics but rather to robustly evaluate relative differences across experimental states. These relative comparisons, such as shifts in dimensionality or space coverage with anesthetic depth, are highly interpretable and biologically meaningful even with moderate sample sizes. Thus, our approach emphasizes the practical utility of low-dimensional analysis for discerning clear network-level changes, without overinterpreting absolute effect sizes beyond the inherent sampling resolution.

Several potential future directions naturally follow from our work. One intriguing avenue would be to leverage the NMF-derived models fit to specific states as "basis sets" for transforming and projecting activity from different states, enabling quantitative comparisons across conditions in a common low-dimensional subspace. This could allow a more nuanced understanding of how network structure shifts with perturbations such as anesthesia, aging, or disease. Moreover, future work could integrate constraints or priors more deeply into the decomposition process to reflect known network motifs or anatomical features, moving toward semi-supervised or guided dimensionality reduction. Additionally, the use of dynamic or adaptive dimensionality, adjusting component number over time in single states, could further improve resolution of state transitions and network remodeling.

While the main analyses in this study focused on spontaneous NMF, fully unsupervised decomposition and dimensionality, we also implemented a constrained version (CNMF) to allow the imposition of specific analytical designs or hypotheses. CNMF has been widely applied to trace extraction and motion correction tasks^24,36–39^. Here, we implement and reframe as a flexible analytical tool for controlled, hypothesis-driven analysis of low-dimensional structure, opening new possibilities for targeted network investigations.

Beyond NMF and CNMF, we also implemented the standard and widely employed methods of PCA, ICA, and UMAP in our toolbox. While NMF is an exceptional method for optimizing for these low dimensional representations of macroscopic activity, each of these dimensionality reduction techniques offers distinct advantages depending on analytical goals: PCA identifies orthogonal axes of maximal variance; ICA seeks statistically independent sources; UMAP preserves both local and global relationships through nonlinear manifold embeddings. By integrating these approaches, we provide end-users flexibility to tailor dimensionality reduction to the specific needs of their biological questions.

In addition to building tools for activity analysis, we, further, incorporated simulation modules that allow users to generate synthetic neuronal network recordings with tunable properties. These modules, implementing both interconnected and independent process models^18^, provide a powerful exploratory tool for validating dimensionality reduction strategies and understanding model behavior under controlled conditions.

Nonetheless, important limitations must be acknowledged. A major constraint in applying dimensionality reduction methods is the necessity of balancing the sample size of animals against the inherent complexity of the data. While dimensionality reduction is predicated on the assumption that k≪n, where k is the number of components and n is the number of neurons, statistical power ultimately derives from the number of independent biological replicates. In practice, acquiring sufficient sample sizes of high-resolution calcium recordings can be labor-intensive and technically demanding. Thus, interpretations drawn from low-dimensional structures must always be made with careful consideration of the biological sampling resolution. In conclusion, here we introduce an accessible, flexible, and extensible framework, wrapped into a software toolbox, for applying dimensionality reduction to neuronal calcium imaging data, validated across multiple species and conditions. Our results demonstrate that standardized dimensionality reductive modeling not only reproduces prior observations but extends them, offering new biological insights and frameworks for interpreting network dynamics. We anticipate that this toolbox will facilitate a broader adoption of systematic dimensionality reduction in neuroscience, enabling researchers to more easily reveal structure within complex neuronal populations and recordings.

## Methods

### Dimensionality reduction (DR) methods

#### Nonnegative Matrix Factorization (NMF)

We focus our analysis on implementing Nonnegative Matrix Factorization (NMF) because of its superior ability to analyze calcium signals^18^. NMF is a linear decomposition DR technique that aims to factor a given matrix into two constitutive lower rank matrices, where all elements of all matrices are constrained to be nonnegative. Given a matrix X ∈ ℝ^m×n^ , NMF aims to decompose X into two lower rank matrices W ∈ ℝ^m×k^ and H ∈ ℝ^k×n^, such that:

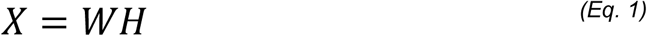

where we can capture a significant portion of the variance at k << min(m,n)^17^, and the multiplication of W and H optimally approximates X for k^23^. The accuracy of the approximation is described by the variance explained as represented by the coefficient of determination (R^2^) ^40^, and is given by:

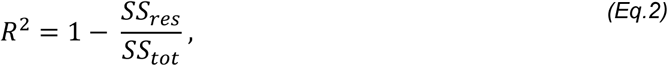

where,

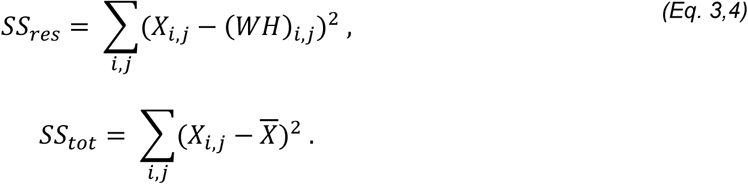

with the model better decomposing the original data in X as R^2^ increases, perfectly capturing it as the limit of the variance explained approaches one. A major advantage to NMF is the adaptation of Akaike information Criterion (AIC) to select the number of dimensions for decomposition ^30,31^, where the optimal number of components minimizes^29^:

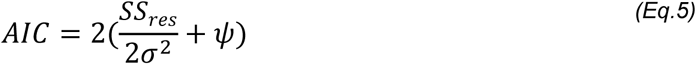

where SS_res_ is the sum of the squared residuals (Eq. 3), σ^2^ is the estimated variance of the data set, and ψ are the free parameters in the model (the number of dimensions being decomposed into k, multiplied by the sum of the dimensionality of X, m + n). AIC provides a more objective metric for optimization, as opposed to the “elbow” method used in other DR contexts^19,41^, directly informing the optimal dimensionality of the reduction.

Linearity lends itself to more interpretable model representations at the cost of going beyond very low dimensionality as is common in non-linear manifold learning^41,42^. We can interpret each entry of each row of H as a neuron’s contribution to that component or sub-network of activity, and each column of W is then interpreted as the time-series for that component. Nonnegativity furthers the interpretability conferred by linearity because the data are modeled as the sum of the components, where each component can be interpreted as a specific combination of input features, representing distinct sources of variation in the data^27^.

#### Modeling NMF low-dimensional component spaces in three-dimensions

Previous research has used linear components and non-linear components modeled in three-dimensional space, found in W, to further analyze the neuronal dynamics^10,13,16,32–35^. We do so here too, and also build an expanded framework for low-dimensional component space analysis.

#### Fitting geometries to component activity in three-dimensions

To quantify the extent of neuronal activity represented in three-dimensional space, we fit geometric closed surfaces to the derived curves. The enclosed volume provides a metric to describe the spatial expanse of neuronal activity. Specifically, we apply standard convex hulls^43^ or alpha shapes^44^, using established computational libraries^45,46^ to define precise geometric boundaries around activity patterns. A convex hull is the smallest convex set enclosing all points, often resulting in an overestimation of volume due to its convexity constraint, while alpha shapes provide a more flexible fit, capturing concave features by adjusting a tunable parameter (alpha) that determines the granularity of the boundary. The resulting volume enclosed by these surfaces provides a quantitative measure to characterize the spatial dynamics and expanse of neuronal activity.

#### Analyzing the movement of component activity in three-dimensions

Following previous research, we quantify the smoothness of activity trajectories by calculating the angles between consecutive segments, where smaller angles represent smoother transitions^32^ indicative of consistent neuronal dynamics. Conversely, larger angles can identify abrupt transitions or instability in neuronal activity. Additionally, we evaluate the dynamics through computed distances, defined as the Euclidean distance between consecutive points along the curve. Higher distance values indicate rapid shifts in neuronal network states or activity bursts, whereas lower distances correspond to periods of relative stability or sustained network states.

#### Constrained Non-Negative Matrix Factorization (CNMF)

Another major advantage of NMF is that it can be constrained after initialization to force optimization towards a more biased or specified model^47^. CNMF has become state of the art for automated trace extraction of calcium recordings^24,36–39^. However, constraining has yet to be implemented for more targeted analysis of dynamics. We do so here.

Similarly to NMF, CNMF satisfies:

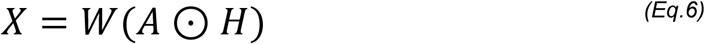

where A is a constraint matrix that enforces constraints on selected entries of H. Specifically, A has the same dimensions as H and contains values indicating which elements of H remain free (1) and which are constrained (0). The elementwise multiplication A ⊙ H then applies these constraints directly, effectively enabling or disabling individual elements of H. We note that for this implementation to remain constrained, multiplicative solvers must be used^47^.

We validate our implementation of CNMF using artificially generated datasets. Specifically, we generate random Gaussian data for X, W, and H, and incorporate constraints through a randomly generated binary matrix A. This ensures that each element of H is properly constrained according to A during the optimization process (Fig. 5). We provide the pseudocode for an implementation of CNMF below:

**Fig 5.**
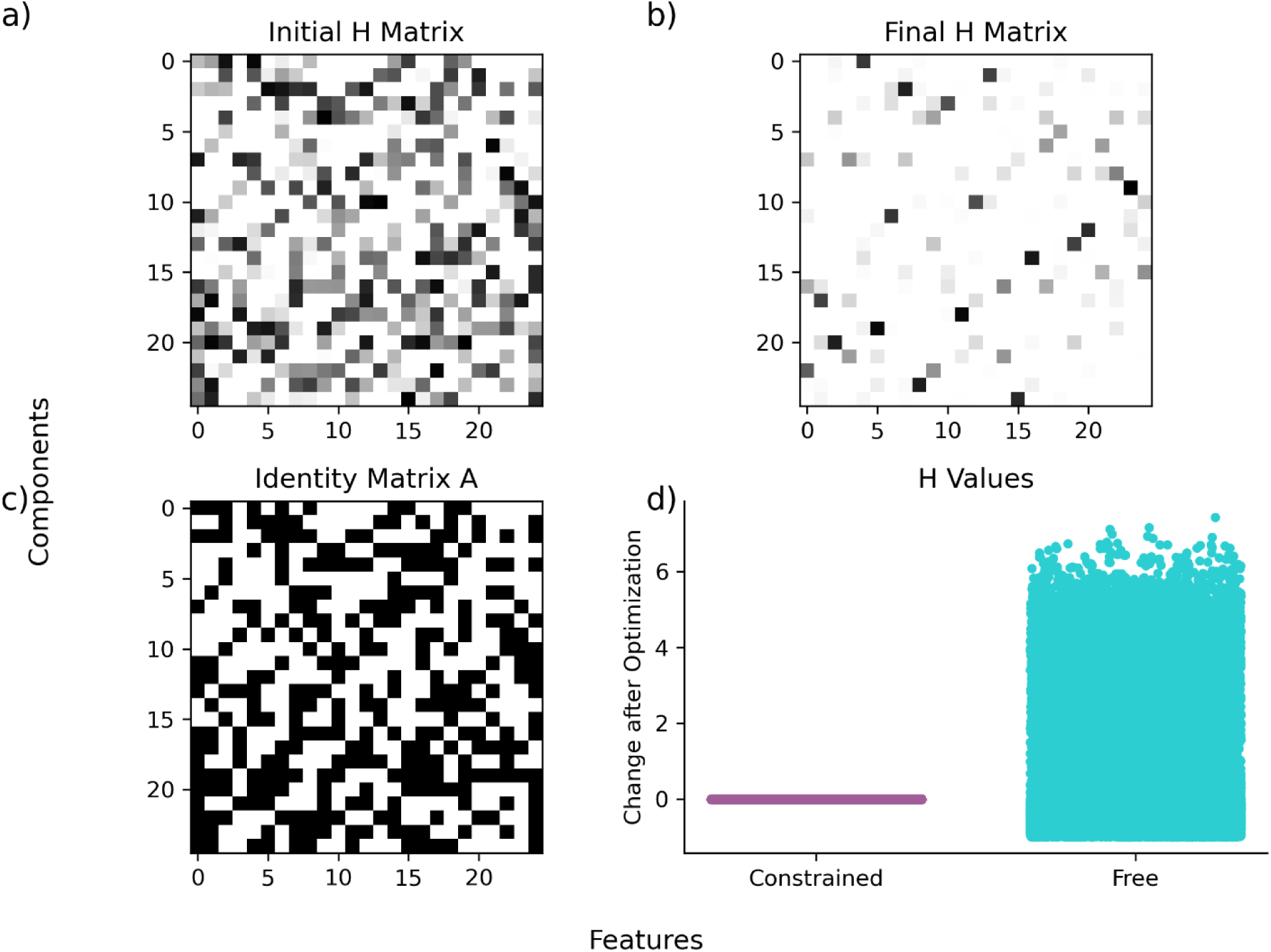
Constrained Non-Negative Matrix Optimization Proof of Constrained Optimization. **a)** Initialized 25x25 random matrix before optimization. **b)** Same 25x25 matrix as shown in a), after optimization. **c)** An initialized binary matrix to constrain the matrix shown in a). **d)** Difference between weights, before and after optimization, split according to constraint status.

Input: X [n_samples x n_features]

1. W, H = init_WH(n_components = target_component_number)
2. A = init_A # this is user defined for analysis purposes
3. model_constrained = optimize_with_constraints(X, W, H, A)

*the only differences between an NMF implementation and the above CNMF implementation are in the necessary initializing of the constraint, and the providing of the constraint matrix to the optimization function

### Other feature implementations in our software wrapper

#### Other NMF initializations

For our analyses, we initialize using nonnegative singular dual value decomposition, a proven method to improve model convergence speed^48^. However, a variety of methods can be used to initialize an NMF with advantages, disadvantages, and different end goals for what components should capture^49,50^. Here we also implement custom, nonnegative PCA, and k-means clustering initialization methods, in addition to the ones provided by sklearn^51^, to allow for more granular user control for analysis, across the general different end goals of initializations^49,50^.

#### Other DR methods

In addition to NMF and CNMF, our software wrapper integrates a comprehensive set of widespread and previously characterized dimensionality reduction (DR) approaches in the context of this analysis^18^. We employ Principal Component Analysis (PCA), Independent Component Analysis (ICA), and Uniform Manifold Approximation and Projection (UMAP). PCA identifies orthogonal axes of maximal variance, ICA seeks statistically independent components in the data, and UMAP applies a nonlinear approach to preserve both local and global structure in the data. We implement these additional methods to broaden the analytical capabilities of the software and offer a high degree of user customization for more specific or targeted analyses.

#### Neuronal network simulations

Previous research has implemented network simulation methods to generate artificial data to validate model performance^18,26^. Here we build a practical and easily tunable way to implement two simulation paradigms, the interconnected networks model in which neurons exhibit user-defined degrees of mutual influence, and the downstream or independent process model, where each neuron (or group of neurons) are primarily driven by separate input streams^18^. While previous research structured a specific set of connection of parameters for validation, our implementation allows full user tunability of parameters across architecture, firing rate, temporal resolution, and connection distributions^18^, in addition to full tunability of calcium signal simulation as well, to account for a broad range of sensor dynamics^18,26,52^.

### Validation analysis of previously collected experimental data

#### Neuronal network dynamics of C. elegans under increasing isoflurane

Using NMF, we investigate the neuronal network dynamics of previously collected C. elegans recordings under increasing concentrations of anesthesia (0% Isoflurane, 4% Isoflurane, and 8% Isoflurane), for ten animals. Briefly, researchers encapsulated C. elegans in a permeable hydrogel and used light-sheet microscopy to image nearly all head neurons (n = 150) expressing GCaMP6s (a calcium indicator) at two volumes per second to measure fluorescent activity^16^. Neurons were tracked over time, across conditions, and normalized activity extracted^16,53–56^.

Neuronal network dynamics of murine layer 2/3 in murine S1 under increasing isoflurane:

We further investigate neuronal network dynamics of layer 2/3 of murine S1, also under increasing concentrations of isoflurane (0%, 0.7%, 1.4%, and recovery), for 36 animals. Previous work recorded two-photon calcium imaging in head-fixed mice at a frame rate of 29.16 Hz^28^, systematically varying the inspired isoflurane concentration. The same neurons were also tracked across conditions^28^, and activity also extracted^24,28,38^.

#### Recording analysis pipeline

We develop and implement a standardized pipeline to analyze both recording sets. For each animal in either data set, we first set the minimum of all traces to 0, to satisfy the constraints of nonnegativity. We then concatenate the recordings across all experimental states to globally normalize activity across each animal, setting the animal minimum to 0 and the maximum to 1. With traces on the same scale across experimental contexts, for each experimental condition, we begin by iteratively fitting models for the first twenty-five components to quantitatively characterize the profile of variance explained and information criterion as a function of adding components. Further, by characterizing the information profile for a particular recording, we can also find the optimal number of components to describe the recording^18^, at the minimum of the AIC curve^29^.

We continue by fitting the recording to the optimal number of components to analyze neuronal contributions (found in H) [Eq. 1] to patterns of activity in an unsupervised learning fashion. Finally, we fit a three-component model to apply NMF in three-dimensional space. Recent DR analysis of our data revealed oscillatory dynamics in three-component space^16^. Here, we extend these previous findings by quantifying the oscillatory properties through standard spectral analysis^57^. We conclude by finding the space encapsulating the activity in three-dimensional space, volume, and characterizing its movement about space, distance and direction.

We provide the pseudocode for our analytical pipeline below:

Input: animal_recordings [time points x neurons x experimental epochs]

1. make trace minimum for all traces across epochs 0
2. concatenate traces [time points * experimental epochs x neurons]
3. normalize traces from (0,1)*
4. split traces into original dimensions [time points x neurons x experimental epochs]
5. for epoch in epochs:
6. initialize list of variance explained, and AICs
7. for i in range(1, num_components) # we use 25
8. W, H = init_WH(n_components = i)
9. NMF_model = optimize(X, W, H)
10. var_ex_list.append(NMF_model.var_ex)
11. AIC_list.append(NMF_model.aic)
12. optimal_components = min(AIC_list)
13. W_optimal, H_optimal = init_WH(n_components = optimal_components)
14. model_optimal = optimize(X, W_optimal, H_optimal)
15. W_state_space, H_state_space = init_WH(n_components = 3)
16. model_state_space = optimize(X, W_state_space, H_state_space)
17. volumes, directions, distances = analyze_state_space(model_state_space.W)

* we normalize using the global maximum and minimum of the trace set to preserve the relative difference between them.

## Acknowledgements

A very warm thanks to Dr. Jacob F. Norman for feedback and guidance in the development of this research. We would also like to thank Dr. Christopher W. Connor for his support in collecting the murine anesthetic data.

## Data and Code Availability Statement

### Author Contributions

Conceptualization, all authors; Methodology, DC; Validation, DC; Formal Analysis, DC; Investigation, DC; Resources, DC, JN, and CVG; Software, DC; Data Curation, DC , JN, and CVG; Writing – Original Draft, DC; Writing – Review and Editing, all authors; Visualization, DC; Supervision, JN, CVG, MAK, and JAW; Project Administration, DC; Funding Acquisition, DC and JAW.

